# Dual role of auxin in regulating plant defense and bacterial virulence gene expression during *Pseudomonas syringae Pto*DC3000 pathogenesis

**DOI:** 10.1101/2019.12.29.881581

**Authors:** Arnaud T. Djami-Tchatchou, Gregory A. Harrison, Chris P. Harper, Renhou Wang, Michael J. Prigge, Mark Estelle, Barbara N. Kunkel

## Abstract

Modification of host hormone biology is a common strategy used by plant pathogens to promote disease. For example, the bacterial pathogen *Pseudomonas syringae* strain *Pto*DC3000 produces the plant hormone auxin (Indole-3-acetic acid, or IAA) to promote *Pto*DC3000 growth in plant tissue. Previous studies suggest that auxin may promote *Pto*DC3000 pathogenesis through multiple mechanisms, including both suppression of salicylic acid (SA)-mediated host defenses and via an unknown mechanism that appears to be independent of SA. To test if host auxin signaling is important during pathogenesis, we took advantage of *Arabidopsis thaliana* lines impaired in either auxin signaling or perception. We found that disruption of auxin signaling in plants expressing an inducible dominant *axr2-1* mutation resulted in decreased bacterial growth, demonstrating that host auxin signaling is required for normal susceptibility to *Pto*DC3000, and this phenotype was dependent on SA-mediated defenses. However, despite exhibiting decreased auxin perception, *tir1 afb1 afb4 afb5* quadruple mutant plants lacking four of the six known auxin co-receptors supported increased levels of bacterial growth. This mutant also exhibited elevated IAA levels, suggesting that the increased IAA in these plants may promote *Pto*DC3000 growth independent of host auxin signaling, perhaps through a direct effect on the pathogen. In support of this, we found that IAA directly impacted the pathogen, by modulating expression of bacterial virulence genes, both in liquid culture and in planta. Thus, in addition to suppressing host defenses, IAA acts as a microbial signaling molecule that regulates bacterial virulence gene expression.

## INTRODUCTION

*Pseudomonas syringae* is a gram-negative bacteria and a causal agent of leaf spot, leaf blight, leaf speck, and bacterial canker disease of tomato, Arabidopsis, and many cultivated crops and ornamental plant species all over the world (1). *P. syringae* is an extracellular hemibiotrophic pathogen which colonizes the surface of host plants as an epiphyte, and later the intercellular space (apoplast) of the infected plant as a pathogenic endophyte. Once in the apoplast, *P. syringae* suppresses basal defense responses by using the Type III protein secretion system (T3SS), that delivers effector proteins directly into host cells. These effector proteins suppress host defenses and presumably alter other aspects of host physiology to elicit the release of nutrients and water from plant cells (2, 3). *P. syringae* then takes up nutrients and water, multiplies to high levels, and causes development of disease. During *P. syringae* pathogenesis, the levels of several plant hormones, including the auxin indole-3-acetic acid (IAA), increase in infected host tissue (4–7).

Although auxin has long been known to be an important virulence factor for gall-forming pathogens and root-associated bacteria (7), it has more recently been discovered to be important during infection by leaf spotting pathogens such as *P. syringae* strains *Pto*DC3000 and *Pma*4326 (4, 8–10). For example, studies by Wang et al. (10) demonstrated that treatment with exogenous auxin suppresses salicylic acid (SA)-mediated defenses in *A. thaliana*. We observed that plants infected with the *Pto*DC3000 *aldA* mutant, which is impaired for IAA synthesis (11), supported reduced growth of the pathogen and that this was correlated with elevated expression of the defense gene *Pathogenesis-related 1* (*PR1*). The reduced growth of the *aldA* mutant was restored to normal levels in *sid2-1* mutant plants, which have impaired SA biosynthesis, suggesting that IAA promotes pathogen virulence by suppressing SA-mediated defenses (11). However, in a separate study we found that elevated auxin levels in transgenic plants overexpressing the *YUCCA1* auxin biosynthesis gene do not promote susceptibility simply by suppressing SA-mediated defenses (8). This suggested that auxin also promotes susceptibility to *Pto*DC3000 by acting independently of SA.

An additional mechanism by which auxin may promote disease susceptibility is by altering host auxin signaling and physiology. Generally, auxin induces transcriptional changes by promoting ubiquitin (Ub)-mediated degradation of AUX/IAA transcriptional repressors. The degradation of AUX/IAAs leads to activation of Auxin Response Factors and expression of auxin responsive genes (12). Previous studies showed that *P. syringae* promotes pathogen growth and disease development through the action of the Type III secreted effector protein AvrRpt2, which interferes with plant auxin signaling by promoting the degradation of AUX/IAA proteins, thereby increasing auxin sensitivity in the host (4, 13). Further, Navarro et al (9) reported that basal defense responses induced against *P. syringae* results in stabilization of the AUX/IAA proteins and down-regulation of auxin-signaling, suggesting that inhibition of auxin signaling may be an important aspect of plant defense. If auxin-induced changes in the host are important during pathogenesis, we predict that host auxin signaling should contribute to host susceptibility, and host auxin responsiveness may be required for suppression of SA-mediated defenses upon infection by *Pto*DC3000.

In addition to a role for auxin in modulating basal host defenses, previous findings suggested that IAA promotes *Pto*DC3000 pathogenesis through one or more mechanisms that function independently of suppression of SA-mediated defenses (8). One possible mechanism is by directly impacting the pathogen and auxin has been shown to act as a signal molecule that can regulate gene expression in plant-associated bacteria (14–18). The objective of this study is to elucidate the role(s) that auxin plays during *Pto*DC3000 pathogenesis. Here we demonstrate that host auxin signaling is required to promote susceptibility to *Pto*DC3000, by suppressing SA mediated defense. Secondly, we show that IAA acts as a microbial signaling molecule that regulates expression of bacterial virulence genes.

## RESULTS

### Plants expressing the dominant *axr2-1* mutation exhibit impaired auxin responses

Modulation of plant hormone physiology is an important virulence strategy for many plant pathogens, and several have been shown to target different aspects of host auxin biology, suggesting a role for auxin signaling in pathogenesis. Consistent with this hypothesis, a previous report by Wang et al. suggested that host auxin signaling may play a role in susceptibility to *P. syringae*, as the *A. thaliana axr2-1* mutant, which is impaired in auxin responses (19), had slightly reduced susceptibility to *P. syringae pv. maculicola* (*Pma*4326) when inoculated with 1×10^6^ cfu ml^-1^ bacteria (10). However, as *axr2-1* mutant plants are developmentally abnormal and severely dwarfed, it is difficult to interpret this observation. The dominant *axr2-1* allele encodes a mutant form of Axr2, an Aux/IAA protein, that is not degraded upon auxin treatment, and therefore auxin responses are not normally induced in this mutant. To further investigate the contribution of host auxin signaling to *P. syringae* pathogenesis, we took advantage of an *A. thaliana* transgenic line expressing a form of the Axr2-1 mutant protein that is translationally fused to the glucocorticoid receptor (20), which allows us to induce nuclear localization of the Axr2-1 upon Dexamethasone (Dex) treatment. Use of this line provides us with plants that grow fairly normally to maturity, and then allows us to disrupt auxin signaling at the time of infection with Dex treatment.

To confirm that expression of the *axr2-1* mutation in these plants disrupts auxin signaling, we treated the adult *GR-axr2-1* transgenic plants with Dex and then monitored auxin responsive gene expression. Mature, 4-5 week old *GR-axr2-1* plants and WT Col-0 plants were sprayed with 10 mM Dex (suspended in 0.1% ethanol) or 0.1% ethanol (mock) 24 hours prior to auxin treatment. Before the Dex treatment, the *GR-axr2-1* transgenic plants looked normal, although they were slightly smaller than WT Col-0 plants. However, one day after Dex treatment the plants exhibited abnormal morphology, in which the younger and newly emerged leaves did not expand normally, and exhibited a down-ward curled shape (Supplemental Fig. 1). Col-0 WT plants treated with Dex and mock-treated *GR-axr2-1* plants did not show this response, suggesting that *axr2-1* impairs auxin-mediated events, such as expansion of young leaves (21).

We monitored expression of the auxin-responsive genes *GH3.3* and *IAA19* in these plants in response to application of 1 µM of the synthetic auxin Naphthaleneacetic acid (NAA) or a control (0.01 % DMSO) using quantitative RT-PCR (RT-qPCR). Transcripts of both *IAA19* and *GH3.3* accumulated to high levels in WT Col-0 plants treated with NAA compared to the control (Fig. 1A-B). In contrast, in plants carrying the *axr2-1* mutation, pre-treated either with Dex or ethanol (mock), the expression of both *GH3.3* and *IAA19* were not induced in response to NAA (Fig. 1A-B). The observation that the *GR-axr2-1* plants exhibited impaired auxin responses even in the absence of Dex treatment suggests that some nuclear localization of the GR-Axr2-1 protein occurs even in the absence of Dex. Our results indicate that plants expressing the *axr2-1* mutation exhibit impaired auxin responses, and thus these plants can be used to investigate if host auxin signaling is involved during pathogenesis by *Pto*DC3000.

**Figure 1.**
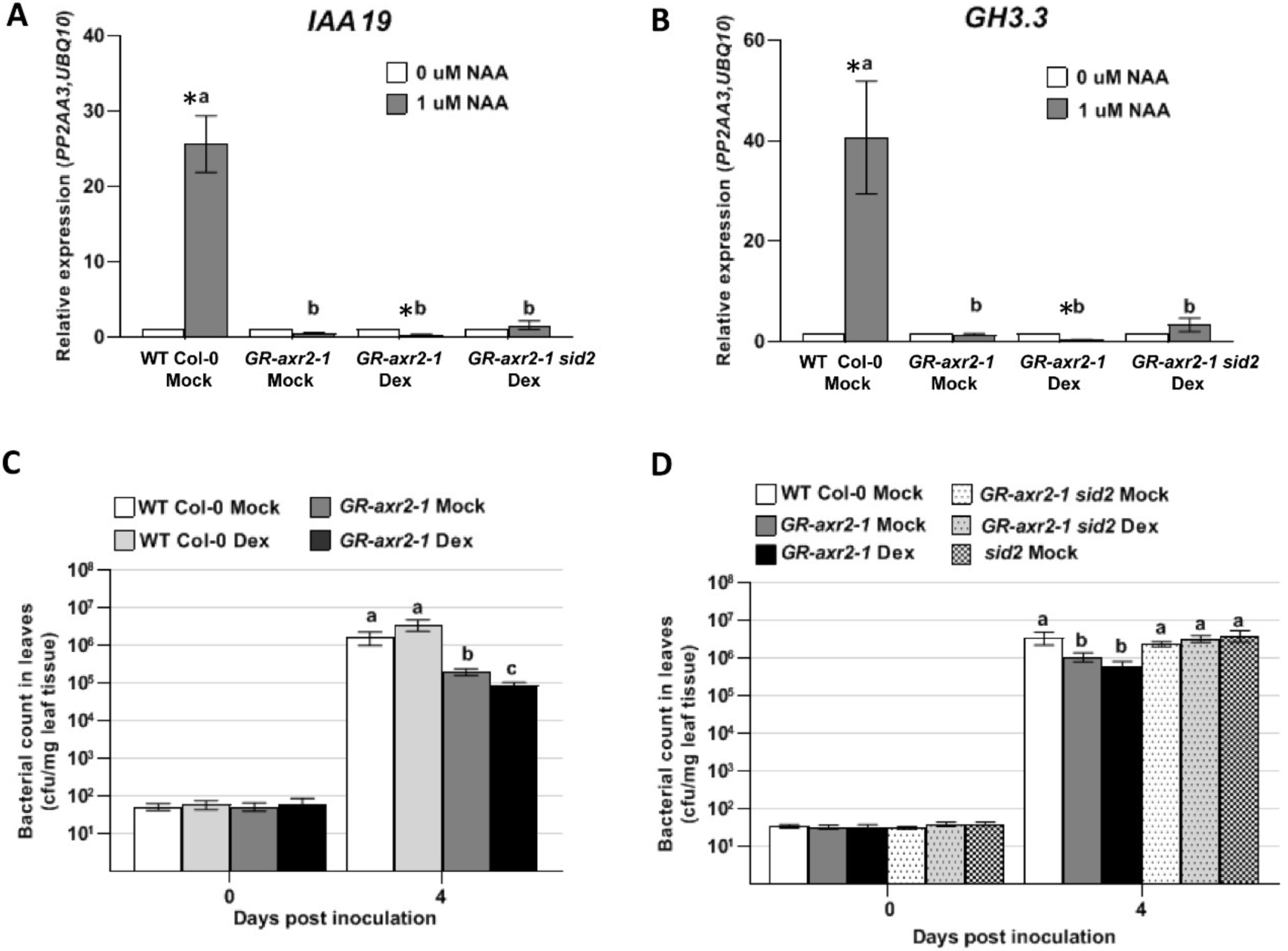
Expression of auxin-responsive genes *IAA19* and *GH3.3* and growth of *Pto*DC3000 in plants carrying the dominant *axr2-1* mutation. Quantitative reverse transcription-polymerase chain reaction (RT-qPCR) analysis of auxin-responsive *IAA19* **(A)** and *GH3.3* **(B)** gene expression 3 hours after treatment with 1 μM Naphthaleneacetic acid (NAA) or 0.01 % DMSO (0 μM NAA). Plants were pre-treated with Dex or a buffer control (Mock) 24 hours prior to NAA treatment. The expression data were normalized using *PROTEIN PHOSPHATASE 2A* SUBUNIT A3 (*PP2AA3*) and *POLYUBIQUITIN* 10 (*UBQ10*) as reference genes and the mock-treated sample was used as a calibrator of relative expression. Values are an average ± standard error of the mean (SEM) for data compiled from 3 independents experiments for Col-0 WT Mock (n= 9) and for 2 independent experiments for *GR-axr2-1* and *GR-axr2-1 sid2* (n= 6). Error bars are too small to see for several data points. **(C)** and **(D)** Quantification of growth of *Pto*DC3000 in WT and GR-*axr2*-1 transgenic plants in plants that were pre-treated with Dex or a buffer control (Mock) 24 hours prior to inoculation. For panel **(C),** values are an average ± SEM for data from 4 independent experiments, carried out on different days, combined to generate composite growth curves, resulting in a total of 12 to 16 biological replicates for day 0 and 20 to 24 replicates for day 4. For panel **(D)**, values are an average ± SEM for data from 3 independent experiments, carried out on different days, combined to generate the composite growth curves, resulting in a total of 12 biological replicates for day 0 and 20 replicates for day 4. Statistical significance between plant genotypes determined by one-way ANOVA, followed by a Tukey’s post test. Samples indicated by different lower case letters are significantly different (p<0.05). * indicates significant difference between treatments (0 μM NAA vs 1 μM NAA) with p<0.05.

### Plants expressing the *axr2-1* mutation exhibit reduced susceptibility to *P. syringae* strain *Pto*DC3000

To study the contribution of host auxin signaling to *Pto*DC3000 pathogenesis, we assayed the *GR-axr2-1* transgenic plants for altered susceptibility to *Pto*DC3000. *Pto*DC3000 grew to high levels in WT Col-0 plants, regardless of whether or not they were treated with Dex (Fig. 1C). In contrast, plants carrying the *axr2-1* mutation reproducibly supported significantly lower levels of *Pto*DC3000 growth compared to the WT Col-0 (Fig. 1C). We also observed that growth of *Pto*DC3000 in the *GR-axr2-1* plants treated with Dex was slightly lower than in mock-treated *GR-axr2-1* plants. Thus, although some GR-Axr2-1 protein appears to enter the nucleus in the absence of Dex, nuclear localization of additional Axr2-1 protein resulted in a further reduction in susceptibility to *Pto*DC3000. These results indicate that host auxin signaling is important for normal susceptibility to *Pto*DC3000.

### Normal disease susceptibility in *GR-axr2-1* plants is restored by introducing the *sid2* mutation

To test if the reduced susceptibility phenotype in *GR-axr2-1* plants is dependent on SA-mediated defenses, we crossed the *salicylic acid induction deficient 2* (*sid2-2*) mutation, which carries a deletion in *ISOCHORISMATE SYNTHASE 1* (*ICS1*) and thus abolishes pathogen-induced SA accumulation (22), into the *GR-axr2-1* transgenic line. Introduction of *sid2-2* did not substantially alter the reduced auxin-responsiveness in plants expressing the *axr2-1* mutation (Fig. 1A-B). However, during infection, the *GR-axr2-1 sid2-2* plants supported similar levels of *Pto*DC3000 as WT Col-0 and *sid2-2* plants (Fig. 1D). Thus, introduction of the *sid2-2* mutation restores normal levels of susceptibility to *Pto*DC3000 in *GR-axr2-1* plants, suggesting that the reduced susceptibility in these plants with impaired auxin-signaling is due to elevated SA-mediated defenses. This is consistent with previous observations suggesting that SA and auxin signaling are mutually antagonistic in *P. syringae*/Arabidopsis interactions (9–11).

### TIR1/AFB auxin co-receptor mutants retain susceptibility to *Pto*DC3000 infection

Given our observation that host auxin signaling contributes to disease susceptibility, we were interested in determining if any specific auxin co-receptors play a role in *Pto*DC3000 infection. The genome of *A. thaliana* encodes six auxin co-receptors (TIR1 and AFB1-5) that mediate diverse responses to the plant hormone auxin (23, 24). We took advantage of existing auxin co-receptor mutants to investigate the contributions of various combinations of the six known TIR1/AFB family proteins during *Pto*DC3000 pathogenesis. We tested two higher order mutants that lack four TIR1/AFB proteins, but are still able to develop into mature plants: the *tir1-1 afb1-3 afb2-3 abf3-4 (tir1 afb1 afb2 abf3)* quadruple mutant (25), and the *tir1-1 afb1-3 afb4-8 afb5-5* (*tir1 afb1 afb4 afb5*) quadruple mutant (26). Although *tir1 afb1 afb2 afb3* mutant plants are severely impaired developmentally as seedlings, and are dwarfed as adult plants, they supported normal levels of *Pto*DC3000 growth, similar to that observed in WT Col-0 (Suppl. Fig. 2A). We also observed that *tir1 afb1 afb4 afb5* mutants did not exhibit reduced disease susceptibility to *Pto*DC3000 (Fig. 2A). Unexpectedly, the *tir1 afb1 afb4 afb5* mutant actually supported significantly higher levels of pathogen growth compared to the WT Col-0 plants (Fig 2A and Suppl. Fig. 2B). These results indicate that inactivation of 4 out of 6 auxin co-receptors does not compromise susceptibility to *Pto*DC3000.

**Figure 2.**
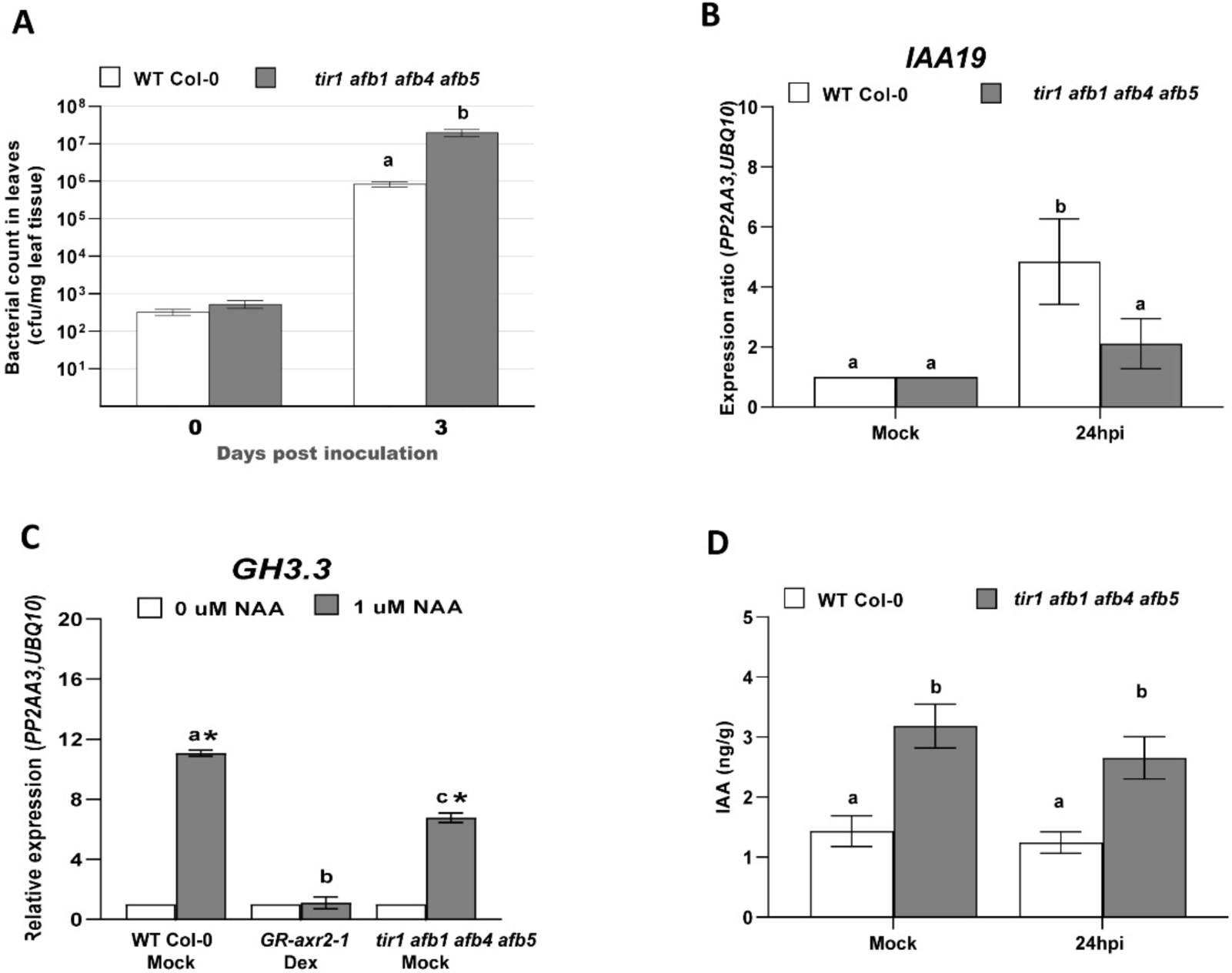
Growth of *Pto*DC3000, IAA-responsive gene expression and quantification of IAA levels in auxin receptor mutants. **(A)** Growth of *Pto*DC3000 in the *tir1 afb1 afb4 afb5* mutant. Values are an average ± SEM for data from 4 independent experiments, carried out on different days, combined to generate composite growth curves, resulting in a total of 4 to 6 biological replicates for day 0 and 8 replicates for day 3. Statistical significance between plant genotypes was analyzed using the student’s t-test. **(B)** Expression of the auxin-responsive gene *IAA19* in WT and *tir1 afb1 afb4 afb5* mutant plants at 24 hours after *Pto*DC3000 inoculation. Values are an average ± SEM for data compiled from 2 independent experiments with 6 biological replicates in total. **(C)** Auxin-responsive *GH3.3* gene expression in WT, *GR-axr2-1* and *tir1 afb1 afb4 afb5* mutant plants 3 hours after treatment with 1 μM NAA or 0.01% DMSO (0 μM NAA). Plants were pre-treated with Dex or a buffer control (Mock) 24 hours prior to NAA treatment. Values are an average ± SEM for 3 biological replicates for Col-0 WT Mock, *GR-axr2-1* and *tir1 afb1 afb4 afb5*. * indicates significant difference between treatment (0 and 1 μM NAA) with p<0.05. Similar results were observed for *tir1 afb1 afb4 afb5* in a 2^nd^ independent experiment. For panels B & C, the expression data were normalized using *PROTEIN PHOSPHATASE 2A* SUBUNIT A3 (*PP2AA3*) and *POLYUBIQUITIN* 10 (*UBQ10*) and the mock-treated sample was used as a calibrator of relative expression. **(D)** Free IAA levels in WT Col-0 and *tir1 afb1 afb4 afb5* mutant plants, 24 hours after Mock treatment (10 mM MgCl_2_) or inoculation with *Pto*DC3000. Values are an average ± SEM for data compiled from 2 independent experiments with 6 biological replicates in total. Results were analyzed using one way ANOVA, followed by a Tukey’s post test. Similar results were obtained in a third independent experiment. Samples indicated by different lower-case letters are significantly different (p<0.05).

We originally expected that these TIR1/AFB co-receptor mutants would exhibit decreased susceptibility to *Pto*DC3000, because we hypothesized that they would have impaired auxin perception and signaling, similar to the *GR-axr2-1* line. However, since the TIR1/AFB co-receptor mutants did not have decreased susceptibility to *Pto*DC3000, we hypothesized that the remaining 2 AFB family members (AFB4 and AFB5 in the *tir1 afb1 afb2 afb3* mutant, and AFB2 and AFB3 in the *tir1 afb1 afb4 afb5* mutant) are sufficient to mediate normal auxin perception and signaling during infection. Therefore, to test if the *tir1 afb1 afb4 afb5* mutant plants exhibit altered auxin perception during *Pto*DC3000 infection, we infected WT Col-0 and *tir1 afb1 afb4 afb5* plants with *Pto*DC3000, and collected total RNA 24 h after inoculation to monitor expression of the auxin-responsive gene *IAA19*. In WT Col-0 infected plants, we found that, as expected, the expression of *IAA19* was induced by 24 hours after infection, consistent with previous data that *Pto*DC3000 infection results in elevated IAA levels (4,5,6). In contrast, expression of *IAA19* was not significantly induced in *tir1 afb1 afb4 afb5* plants at 24 hours after infection (Fig. 2B), suggesting that this mutant exhibits reduced auxin perception and/or signaling during *Pto*DC3000 infection.

However, since the *tir1 afb1 afb4 afb5* mutant still harbors intact AFB2 and AFB3 auxin co-receptors, we predicted that the *tir1 afb1 afb4 afb5* mutant plants may not be as severely compromised in auxin signaling as the *GR-axr2-1* plants. To directly compare the auxin responsiveness of these two lines, we treated them with NAA and monitored expression of *GH3.3* and *IAA19*. As observed previously, *GH3.3* and *IAA19* were not induced by NAA in *GR-axr2-1* plants (Fig 2C and Suppl. Fig. 2C). However, both auxin-responsive genes were induced by NAA treatment in the *tir1 afb1 afb4 afb5* mutant, although the level of induction was lower than observed in WT. Thus, auxin signaling in the *tir1 afb1 afb4 afb5* mutant is only partially impaired.

### The *tir1 afb1 afb4 afb5* mutant accumulates elevated levels of IAA

It is not immediately obvious why the *tir1 afb1 afb4 afb5* mutant, which has decreased auxin-responsive gene expression, exhibits enhanced susceptibility to *Pto*DC3000. We hypothesized that the *tir1 afb1 afb4 afb5* plants might accumulate elevated levels of IAA, due to disruption of the feedback mechanism that normally maintains IAA homeostasis in WT plants (27) and that this elevated IAA promotes susceptibility. To investigate whether the enhanced pathogen growth in *tir1 afb1 afb4 afb5* plants is correlated with elevated IAA in host tissue, we quantified free IAA levels in WT Col-0 and *tir1 afb1 afb4 afb5*, in both *Pto*DC3000-infected and mock-treated plants at 24h post inoculation. We found that *tir1 afb1 afb4 afb5* mutant plants accumulated significantly higher levels of IAA than WT Col-0 plants, regardless of treatment (Fig. 2D). Thus, the elevated levels of IAA in this mutant may promote pathogen growth despite a partial impairment in auxin responsiveness.

### Elevated IAA in *tir1 afb1 afb4 afb5* plants is associated with reduced SA-mediated defenses

Observations from several previous experiments suggest that stimulation of host auxin signaling promotes virulence by suppressing SA-mediated defenses (10, 11). Since *tir1 afb1 afb4 afb5* plants are still capable of responding to NAA, we hypothesized that the elevated levels of IAA in *tir1 afb1 afb4 afb5* plants may promote pathogen growth by suppressing SA-mediated defenses through host auxin signaling via AFB2 and AFB3. Thus, we predicted that induction of SA-mediated defenses would be reduced in these plants compared to in wild-type plants. We tested this by monitoring both the accumulation of SA and the expression of the SA-responsive defense gene *PR1* in infected plants. Although we observed an increase in SA levels in both WT Col-0 and *tir1 afb1 afb4 afb5* plants after inoculation with *Pto*DC3000, the level of SA that accumulated in the mutant was significantly lower than in WT Col-0 (Fig. 3A). Furthermore, *PR1* expression was strongly and reproducibly induced by 24 h post inoculation in WT Col-0 plants, but was not significantly induced over levels observed in mock-treated *tir1 afb1 afb4 afb5* plants (Fig. 3B). Thus, the modest reduction in SA levels in infected *tir1 afb1 afb4 afb5* mutant plants is correlated with the lower expression of *PR1* observed in these plants. These results are consistent with the hypothesis that elevated IAA levels suppress SA-mediated defenses during *Pto*DC3000 infection.

**Figure 3.**
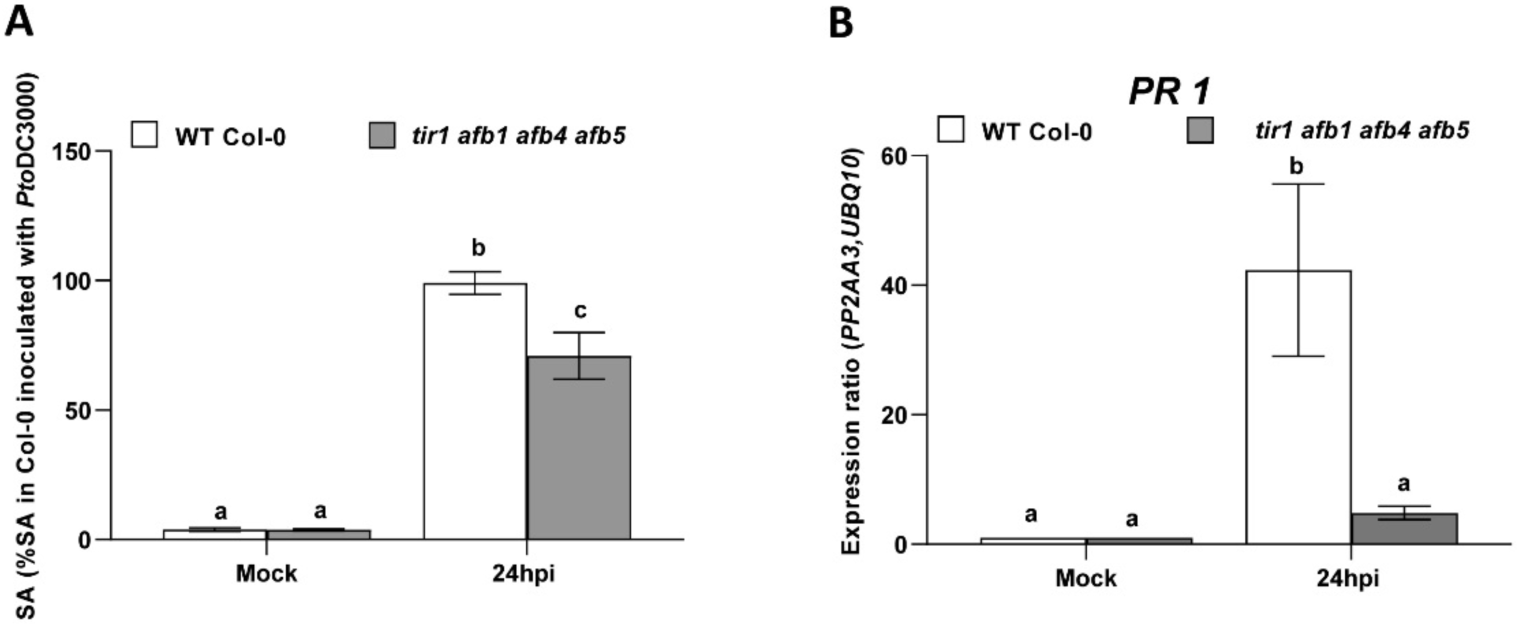
SA-mediated defenses in WT Col-0 and the *tir1 afb1 afb4 afb5* auxin receptor mutant. **(A)** SA levels in WT Col-0 and *tir1 afb1 afb4 afb5* mutant plants, 24 hours after mock treatment (10 mM MgCl_2_) or inoculation with *Pto*DC3000. Values are expressed as a percent of SA accumulation in WT Col-0 24 hours after inoculation with *Pto*DC3000, and represent an average ± SEM for data compiled from 2 independent experiments, with 6 biological replicates per plant genotype per treatment. Results were analyzed using ANOVA, followed by a Tukey’s post test, and different letters indicate significant difference between samples with p<0.05. **(B)** Expression of *PATHOGENESIS RELATED 1 (PR1)* 24 hours after mock treatment (10 mM MgCl_2_) or inoculation with *Pto*DC3000. Expression data were normalized using *PP2AA3* and *UBQ10* to give the relative gene expression. Values are an average ± SEM for data compiled from 2 independent experiments with 6 biological replicates in total.

However, suppression of SA-mediated defenses may not be the only mechanism by which IAA enhances disease susceptibility. Previous findings suggest that IAA can promote *Pto*DC3000 pathogenesis through one or more mechanisms that function independently of SA-mediated defenses (8). Furthermore, our data show that auxin-responsive gene expression is not significantly induced in *tir1 afb1 afb4 afb5* plants during infection (Fig. 2B), suggesting that this mutant does not exhibit significantly increased auxin responses despite elevated IAA levels. Therefore we reasoned that the increased bacterial growth in the *tir1 afb1 afb4 afb5* mutant plants may not be mediated solely through host auxin responses, and considered the possibility that the elevated IAA may promote pathogen growth via an additional mechanism.

### IAA regulates *Pto*DC3000 virulence gene expression in culture

One possible mechanism by which IAA can promote pathogen growth independently of host auxin signaling is by directly impacting the pathogen, for example by modulating virulence gene expression, as has been previously shown in *Agrobacterium tumefaciens* (28, 29), *Dickeya didantii* (formerly known as *Erwinia chrysanthemi*; 18) and *P. savastanoi pv. savastanoi* (14). Accordingly, we sought to determine if IAA also directly impacts *Pto*DC3000 virulence gene expression in culture. First, to asses if IAA has an effect on bacterial growth, we cultured *Pto*DC3000 in rich media (NYG) for several hours before transferring cells to *hrp/hrc* de-repressing media (HDM), a minimal media believed to mimic growth conditions in the apoplastic space of the leaf (30, 31), containing a variety of concentrations of IAA (Supplemental Fig. 3A). We observed that transfer of cells to HDM containing 100 μM IAA or higher impaired the growth of the cells, compared to growth in HDM lacking IAA [Supplemental Fig. 3A, 32]. Thus, IAA does impact *Pto*DC3000 biology. We decided to analyze gene expression in cells treated with 100 μM IAA, as this concentration clearly affected *Pto*DC3000.

Central to the pathogenesis of *Pto*DC3000 is the deployment of a Type III secretion (T3S) system (33, 34). Expression of genes involved in T3S (e.g. *avrPto*, and *hrpL*, which encode a T3S-effector and the RNA polymerase sigma factor responsible for transcribing T3S-related genes, respectively), can be induced in culture by growing cells in HDM (30, 31). To test the effect of IAA on expression of virulence genes, we transferred *Pto*DC3000 from NYG to HDM containing 0 or 100 μM IAA, grew the cells for 90 min (Supplemental Fig. 3B) and monitored the expression of both known and putative virulence genes. The housekeeping genes *gyrB* and *rpoD* were used as an internal reference, and gene expression levels were normalized to the basal expression levels in NYG. As expected, we observed low levels of expression of the T3S genes *hrpL* and *avrPto* in NYG, and strong induction upon transfer to HDM (Fig. 4A, B). However, when the cultures were treated with IAA, the induction of *hrpL* and *avrPto* was significantly reduced (Fig. 4A, B). In contrast, the presence of IAA did not alter the expression of *cmaA,* a gene required for synthesis of the virulence factor coronatine (Fig. 4C), and stimulated expression of the virulence-associated gene *tvrR* (35; Fig. 4D) as well as genes predicted to be involved in Type VI secretion (T6S), *hcp1* and *PSPTO_5415* (Fig. 4E, F). We chose to examine these last two genes as T6S systems have been proposed to play a role during bacteria-host interactions, or in bacteria-bacteria interactions in the microbial community. Further, expression of T6S-related genes has been previously shown to be regulated by IAA (14). These results indicate that exposure to IAA causes transcriptional changes in *Pto*DC3000, leading to both up- and down-regulation of distinct subsets of genes.

**Figure 4.**
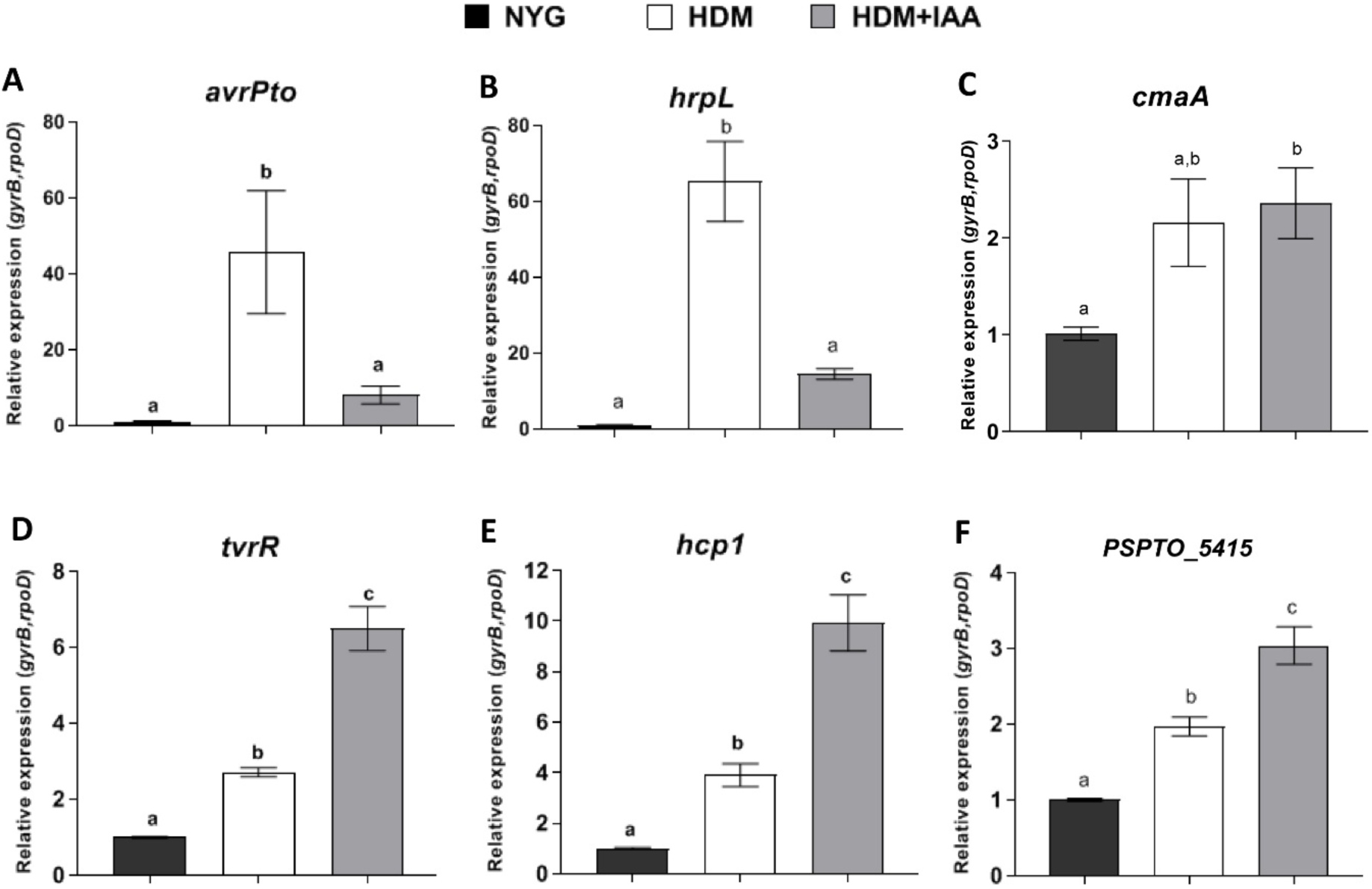
The effect of IAA treatment on *Pto*DC3000 virulence-related gene expression in culture. Expression of virulence-related genes **(A)** *avrPto*, **(B)** *hrpL*, **(C)** *cmaA*, **(D)** *tvrR*, **(E)** *hcp1* and **(F)** *PSPTO_5415* 1.5 hours after being transferred from NYG to HDM or HDM containing 100 μM IAA (HDM+IAA). Expression levels in cells growing in NYG was used as calibrator of relative expression. The relative expression was calculated using the reference genes *rpoD* and *gyrB*. Data from 2 independent experiments, carried out on different days, were combined, with each data point representing the average of 6 biological replicates, and error bars representing the SEM between biological replicates. Results were analyzed using ANOVA, followed by a Tukey’s post test. Different letters indicate significant difference between samples with p<0.05. This experiment was repeated a third time with similar results.

### Auxin modulates bacterial virulence-related gene expression *in planta*

Our results indicate that IAA induces transcriptional changes in *Pto*DC3000 in culture. However, studies performed on bacteria grown in culture may not fully represent bacteria growing in host tissue during infection. Therefore, we next sought to investigate if IAA also impacts *Pto*DC3000 gene expression when growing in plant tissue. To accomplish this, we took advantage of the *tir1 afb1 afb4 afb5* mutant described above, as it accumulates elevated IAA (Fig. 2D). Wild-type Col-0 and *tir1 afb1 afb4 afb5* mutant plants were inoculated with *Pto*DC3000 and leaves were collected at 24 and 48 h post inoculation (Supplemental Fig. 2B). Total RNA, including both *Pto*DC3000 and plant RNA, was purified from these leaves and used to monitor expression of the bacterial virulence-associated genes assayed in culture. RNA from the inoculum used to infect these plants was used as a calibrator of relative expression.

We assessed the stability of expression of several house-keeping genes in order to choose those suitable for use as reference genes for normalization of gene expression (36) and found that *recA* and *rpoD* were expressed at very similar levels at 24 and 48 hours after inoculation. Similar to what has been previously observed, the expression of *avrPto* and *hrpL* was strongly induced within 24 hours after inoculation, compared to the expression level in the inoculum (Fig. 5A, B) (37–39), and the expression of both genes declined by 48 hpi. However, the expression of *avrPto* and *hrpL* was significantly lower in *tir1 afb1 afb4 afb5* mutant plants compared to WT Col-0 plants, a finding that is consistent with our observation that IAA suppresses the induction of these genes in culture (Fig. 4A, B). Thus, elevated IAA also appears to suppress expression of T3S genes during growth in plant tissue. We observed that expression of *cmaA* also was induced in planta, but consistent with our findings in liquid media, its expression was not significantly different in WT and *tir1 afb1 afb4 afb5* plants (Fig. 5C). Furthermore, the expression of *tvrR* was induced in planta and was expressed at higher levels in *tir1 afb1 afb4 afb5* mutant plants, with elevated expression most pronounced at 48 hours after inoculation (Fig. 5D). Interestingly, we did not observe consistent up regulation of *hcp1* and *PSPTO_5415* in these experiments, suggesting that there are some effects of IAA in liquid culture that are not detectable in our experiments in planta. Overall, our results demonstrate that IAA regulates the expression of virulence-related genes in *Pto*DC3000, both in culture and during growth in planta.

**Figure 5.**
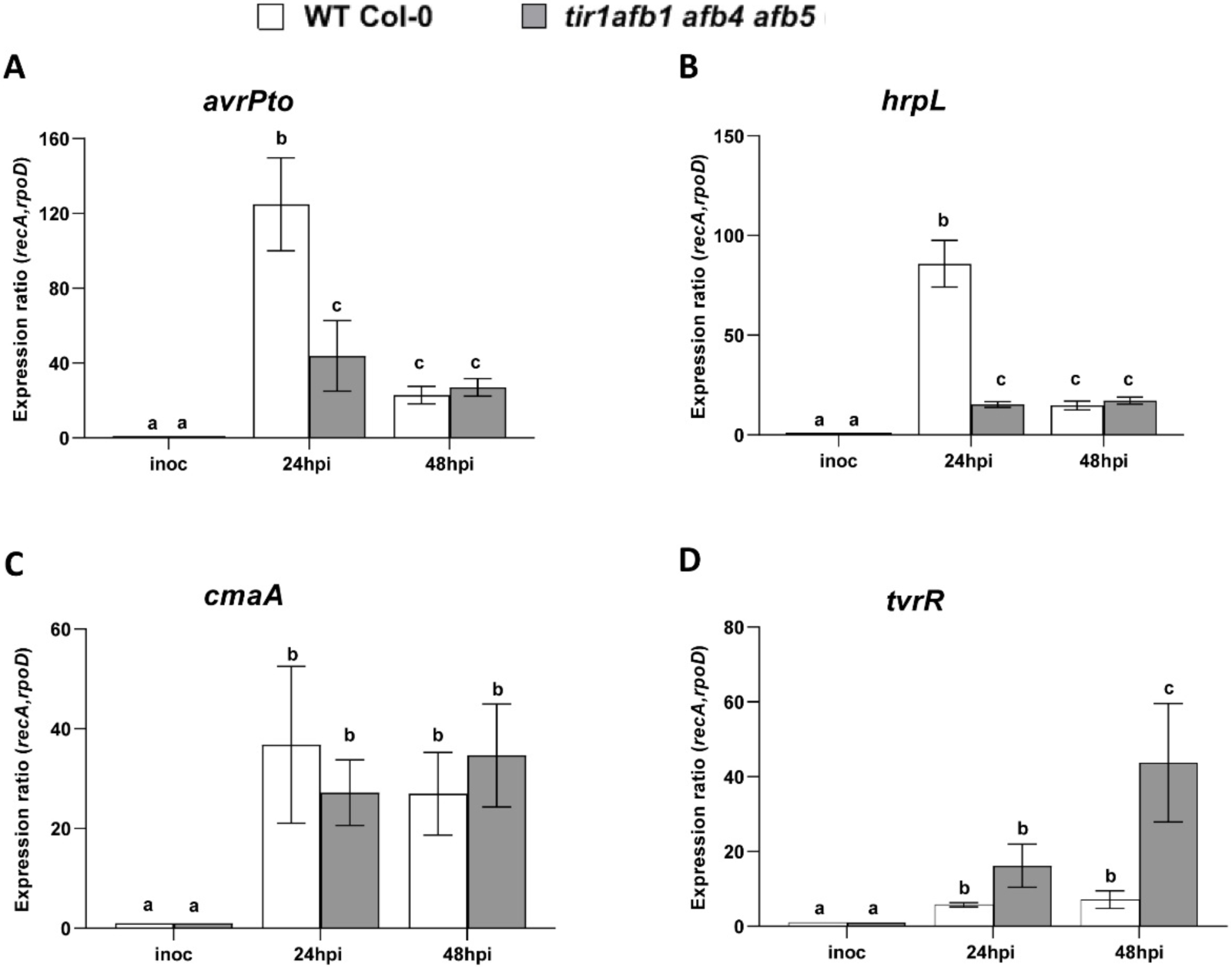
The effect of elevated IAA levels on *Pto*DC3000 virulence-related gene expression *in planta*. Expression of virulence-related genes **(A)** *avrPto*, **B)** *hrpL*, **C)** *cmaA* and **D)** *tvrR* in *Pto*DC3000 growing in WT Col-0 and *tir1 afb1 afb4 afb5 A. thaliana* plants. Infected leaves were harvested 24 and 48 hours after inoculation, and total mRNA isolated and used for RT-qPCR to quantify bacterial gene expression. RNA prepared from the *Pto*DC3000 cell suspension used for the inoculum was used as the calibrator for relative expression. The data shown were compiled from 2 independent experiments, carried out on different days. The relative expression was calculated using the reference genes *recA* and *rpoD*. Each data point is the average of 6 biological replicates, and error bars represent the SEM between biological replicates. Results were analyzed using ANOVA, followed by a Tukey’s post test. Different letters indicate significant difference between samples with p<0.05.

## DISCUSSION

Our investigation of the roles of auxin during pathogenesis of *A. thaliana* by *P. syringae* reveals that auxin promotes virulence of *Pto*DC3000 through two different mechanisms: 1) activating host auxin signaling to suppress SA-mediated plant defenses and 2) directly impacting the pathogen by modulating virulence gene expression.

### Host auxin signaling is required for normal susceptibility to *Pto*DC3000

There is growing evidence that auxin promotes disease development in *A. thaliana*, and it is hypothesized this is mediated primarily through regulatory cross talk between auxin and SA-mediated signaling pathways in the host (11, 40–42). Our finding that an *A. thaliana* transgenic line carrying the dominant *axr2-1* mutation (*GR-axr2-1*) that impairs auxin responses exhibits reduced susceptibility to *Pto*DC3000 is consistent with earlier observations by Wang et al. (10), and confirms that host auxin signaling is required for normal susceptibility to *Pto*DC3000. To further test the hypothesis that auxin signaling promotes disease by suppressing SA-mediated defenses, we introduced the *sid2-2* mutation, and quantified susceptibility to *Pto*DC3000 in these plants. Our finding that wild-type levels of disease susceptibility were restored in *GR-axr2-1 sid2-2* plants (Fig. 1D) indicates that reduced susceptibility in plants with impaired auxin-signaling is due, at least in part, to elevated SA-mediated defenses. This supports the hypothesis that host auxin signaling suppresses SA-mediated defenses in *P. syringae*/Arabidopsis interactions and demonstrates that this is mediated through the canonical TIR1/AUX/IAA host auxin signaling pathway (Fig. 6).

**Figure 6.**
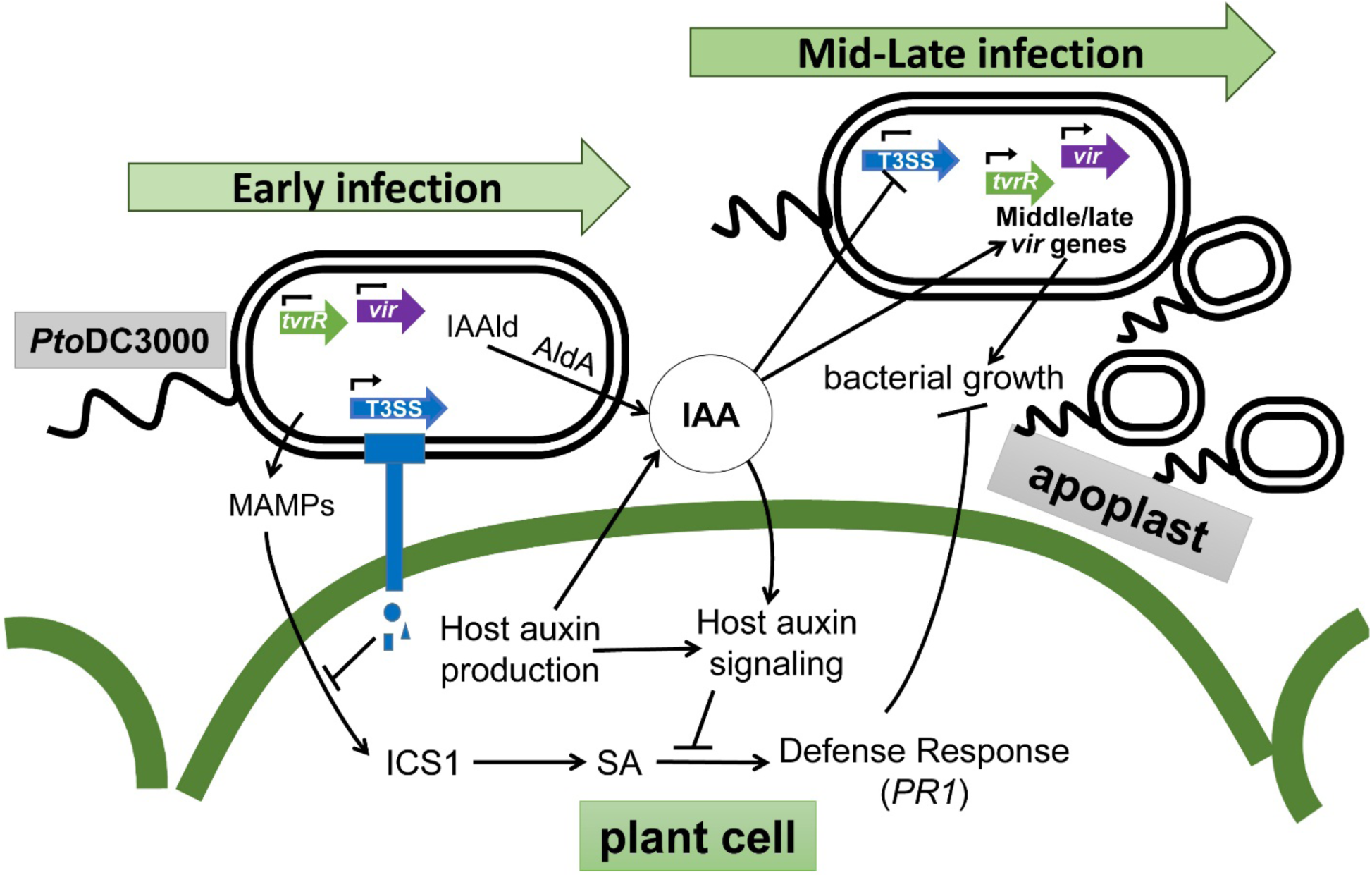
IAA plays multiple roles during *Pto*DC3000 pathogenesis. A working model illustrating how IAA promotes *Pto*DC3000 pathogenesis via multiple mechanisms. Upon *Pto*DC3000 infection, detection of Microbe Associated Molecular patterns (MAMPs) induces expression of basal host defense responses mediated by salicylic acid (SA). Early during infection expression of the Type III Secretion System (T3SS) allows delivery of effector proteins (blue shapes) into the host cell to suppress MAMP-induced defenses. *Pto*DC3000 infection results in elevated auxin (e.g. IAA) levels in infected tissue, possibly due to auxin synthesis by both the host and *Pto*DC3000 (via activity of the Indoleacetaldehyde dehydrogenase AldA, (11). Disruption of host auxin signaling, for example by the dominant *axr2*-1 mutation, prevents suppression of SA defenses, and results in reduced disease susceptibility. IAA also promotes the growth of *Pto*DC3000 independently of suppression of SA-mediated defenses by regulating expression of pathogen virulence genes (green and purple block arrows). We hypothesize that IAA down-regulates T3SS genes after they are no longer needed (e.g. 24 hours post infection), and activates virulence genes such as *tvrR*, that are required at intermediate or late stages of infection. Genes that are transcribed are indicated by the small black arrowhead above the block arrows.

### The *tir1 afb1 afb4 afb5* auxin co-receptor mutant exhibits elevated IAA levels and reduced SA-mediate defenses

The finding that host auxin signaling contributes to disease susceptibility raises the question of whether any specific auxin co-receptors are required for normal *Pto*DC3000 infection. We examined this by testing two higher order (quadruple) *tir1/afb* mutants for altered susceptibility and observed that both mutants supported at least wild-type levels of pathogen growth (Fig. 2 and Suppl. Fig. 2). Although we have not tested all possible combinations of auxin co-receptor quadruple mutants, our observations suggest that the combined activity of two co-receptors (AFB2 and ABF3 in *tir1 afb1 afb4 afb5,* or AFB4 and AFB5 in *tir1 afb1 afb2 abf3*) provides sufficient auxin signaling to support normal levels of pathogen growth. In keeping with this, we demonstrated that while the *tir1 afb1 afb4 afb5* mutant exhibited decreased auxin-responsive gene expression during infection (Fig. 2B), it is only partially impaired in its ability to respond to the synthetic auxin NAA, likely due to the presence of the intact AFB2 and AFB3 auxin co-receptors (Fig. 2C and Suppl Fig. 2C).

Given that host auxin signaling is required for normal pathogenesis, we were surprised to find that the *tir1 afb1 afb4 afb5* mutant exhibited enhanced susceptibility to *Pto*DC3000 (Fig. 2A, Suppl. Fig 2B). The fact that these plants accumulate elevated levels of IAA compared to WT Col-0 (Fig. 2D), provides a reasonable explanation for this result, as we have previously shown that elevated levels of endogenous auxin promotes growth of *Pto*DC3000 (32). Further, our observation that *tir1 afb1 afb4 afb5* plants exhibit elevated IAA levels is consistent with recent findings that homeostasis of endogenous IAA is maintained by feedback regulation through the canonical auxin signaling pathway (27). Consistent with the role for IAA in suppressing SA signaling, we demonstrated that the *tir1 afb1 afb4 afb5* mutant plants exhibited significantly reduced expression of *PR-1* and reduced SA levels compared to WT plants at 24h post inoculation (Fig. 3). These results suggest that during *Pto*DC3000 infection the elevated levels of IAA in the *tir1 afb1 afb4 afb5* plants leads to suppression of SA-mediated defenses. However, it is not clear whether this is mediated primarily at the level of SA synthesis, accumulation, or downstream signaling responsiveness.

### IAA has a direct effect on *Pto*DC3000 by modulating bacterial virulence-related gene expression

Our observation that elevated IAA and increased disease susceptibility in the *tir1 afb1 afb4 afb5* auxin co-receptor mutant is correlated with suppression of SA-mediated defenses conflicts with results from an earlier study, in which we demonstrated that elevated endogenous IAA levels in *YUCCA1* overexpressing plants promoted growth of *Pto*DC3000 via a mechanism independent of SA suppression (8). This discrepancy raised the possibility that auxin also promotes pathogenesis by having a direct effect on the pathogen, for example by modulating virulence-related gene expression. We investigated this by testing the effect of IAA on *Pto*DC3000 gene expression in culture. We observed a significant reduction of *hrpL* and *avrPto* transcript levels when the cultures were treated with IAA (Fig. 4). Initially, we found this result surprising, as we expected that, if IAA impacted virulence gene expression, it would be to promote expression of these genes, which encode virulence factors that are important during early stages of tissue colonization (2, 43). However, a similar negative effect of IAA on expression of T3SS-related genes has recently been reported in the gall-forming pathogen *P. savastanoi* (14). We also observed that IAA did not alter the expression of *cmaA* (Fig. 4C), a coronatine biosynthetic gene which contributes to *Pto*DC3000 virulence (44), and enhanced expression of several other virulence-associated genes, including *tvrR* and T6S-related genes (*hcp1* and *PSPTO_5415,* Fig. 4D-F). Thus, IAA does not to cause a global shift in transcription, but rather differentially impacts specific classes of genes.

IAA has been previously shown to influence gene expression in a variety of plant-associated microbes (7, 16, 45, 46), but this had been only demonstrated in culture, and thus the biological relevance of these findings was unclear. To investigate whether IAA regulates *Pto*DC3000 gene expression *in planta*, we monitored transcript levels of several virulence-related genes in *Pto*DC3000 growing in plants with normal (WT Col-0) or elevated levels of IAA (*tir1 afb1 afb4 afb5* mutants) at 24 and 48 hours post inoculation. As expected, expression of the virulence-related genes examined was induced in both plant genotypes by 24h post inoculation (Fig. 5; 37-39). However, consistent with what we observed in the presence of IAA in culture, induction of *avrPto* and *hrpL* was significantly lower in the *tir1 afb1 afb4 afb5* mutant compared to in WT Col-0 plants at 24h (Fig. 5A,B). Transcript levels of *cmaA* were not significantly different between the two genotypes at either time point, and expression of *tvrR* was elevated in *tir1 afb1 afb4 afb5,* but only at 48 hours post inoculation. Thus, the pattern of IAA-mediated repression or induction of these genes observed in culture reflects the effect of elevated IAA on bacterial virulence genes *in planta*.

Our discovery that IAA appears to directly impact *Pto*DC3000 virulence-related gene expression during growth in *planta* is significant, as it allows us to begin to place the differential regulation of these genes into the context of pathogenesis. Further, it prompts us to hypothesize that IAA acts as a signaling molecule that coordinates expression of virulence genes required during different phases of pathogenesis (Fig. 6). Early during pathogenesis, bacteria colonizing the apoplast assemble the T3SS and secrete T3 effector proteins into host cells in order to suppress basal host defenses (2, 47). Once that is accomplished, the bacteria obtain water and nutrients and multiply to high levels in the apoplast. At this point, the majority of the bacterial cells are not in direct contact with plant cells (48), and presumably do not need to express the T3SS-related genes. By this stage in pathogenesis we also speculate that local IAA levels in the infected tissue have increased to a concentration high enough to down-regulate the T3S-related genes and induce expression of virulence genes involved in subsequent stages of infection (“middle or late virulence genes”, Fig.6). An example of such a gene is t*vrR*, which encodes a transcription factor previously shown to be required for *Pto*DC3000 virulence on *A. thaliana* (35), that we hypothesize regulates transcription of genes required at intermediate stages of infection. Recent experiments by McAtee et al, examining the expression of *P. syringae* pv. actinidiae during infection of Kiwi plantlets reveal three distinct phases of gene expression over the course of infection (37). Their observation that genes encoding the T3SS or effectors are strongly induced during the first 24 hours after infection, and then their expression levels decline by 48 hpi is consistent with what we observed in our studies, and suggests that there is a regulatory mechanism coupling virulence gene expression to the stage of pathogenesis. Future analysis of *Pto*DC3000 virulence gene expression during the various phases of colonization and growth in susceptible host tissue will provide more insight into this fascinating but as yet poorly understood process.

It is now well established that auxin promotes disease development in many plant– pathogen interactions, and in several cases this has been shown to involve suppression of host defenses (40, 11, 42, 49). There is also a growing number of reports that auxin, specifically IAA, has a direct impact on gene expression in plant-associated bacteria (16, 50, 51), and it appears that this regulation plays a variety of different roles. In some interactions IAA may serve as a signal to the microbe that it is in the presence of a potential plant host, and that virulence genes should be induced (7, 18). In others, IAA may serve as a signaling molecule to coordinate gene expression in the microbe, either to induce catabolic genes required for breakdown of IAA (50, 51, 52), or as a signal that a specific stage of infection or a minimal cell density has been achieved, and thus that it is time to activate a new set of genes (53). It is also possible that, in the context of the larger microbial community of the phyllosphere and/or rhizosphere, IAA acts as a microbial signal to regulate interactions between different members of the community (53, 54), and may provide an explanation for the induction of the T6SS by IAA (17, 14). Future studies to investigate the mechanisms by which microbes recognize and respond to IAA, as well as the generation of mutants that do not properly respond to IAA, will allow us and others to more fully examine the multiple roles that auxin and auxin signaling play during plant-microbe interactions.

## MATERIALS AND METHODS

### Plant material and growth conditions

All Arabidopsis thaliana wild type (WT), mutant and transgenic lines used in this study were in the Columbia (Col-0) background. The *sid2-2* mutant was obtained from Mary Wildermuth (22). The *tir1* and *afb1, afb2, afb3, afb4* and *afb5* auxin receptor mutants used in this study, including various higher order mutants, have been previously described (24, 25,26). As the *tir1-1 afb1-3 afb2-3 abf3-4* quadruple mutant displays severe developmental abnormalities, including a high percentage of rootless seedlings or seeds that do not germinate (25), we germinated the mutant on agar plates, transplanted seedlings that produced roots to soil, and allowed them to grow to maturity for pathogen inoculation. Plants were grown on soil in a growth chamber with a short-day photoperiod (8h light/16h dark) at 21°C and 75% relative humidity, with a light intensity of ∼ 130 μ Einsteins sec^-1^ m^-2^.

The *GR-axr2*-1 lines were generated as followed: the DNA fragment encoding the fusion protein *GR-axr2-*1 was made by using fusion PCR. An 834 bp cDNA fragment encoding a rat glucocorticoid receptor was amplified with primers TopoGW_GR_F1 and GR_AXR2_R1, using the pINDEX3 vectors (20) as a template, and *axr2-1* cDNA was amplified with primers GR_AXR2_F1 and AXR2_V1_R1 from cDNA synthesized using RNA extracted from *axr2-1* mutant plants as a template. Purified DNA fragments from these two PCRs were mixed at equal molar ratio and were used as templates and primers for a third PCR to amplify the fusion cDNA *GR-axr2-1*. The fusion cDNA was introduced first into the Invitrogen pENTR/D-Topo vector and then subcloned into the expression vector pEarleyGate 100 (55) downstream of a CaMV35S promoter. The resulting construct (pEarleyGate 100-GR-*axr2-1*) was transferred into Agrobacterium strain GV3101 (pMP90) using electroporation. Transgenic Arabidopsis plants were generated using a floral-dip approach (56).

The *sid2-2 GR-axr2*-*1* line was generated by crossing the *sid2-2* mutant to the *GR*-*axr2-1* transgenic line. F1 progeny from the cross were allowed to self-pollinate, and F2 plants homozygous for the *sid2-2* and the *GR-axr2-1* transgene were identified by PCR genotyping, using primers described in Supplementary Table 1. Plants homozygous for the *sid2-2* allele only yielded an amplification product of 581 bp, indicative of the presence of the *sid2-2* allele (8). Homozygous *sid2-2* plants that scored positively for the presence of the *GR-axr2-1* construct were then allowed to self-pollinate and assayed for segregation of the *GR-axr2-1* construct in the F3 generation. F3 families that segregated 100% for the presence of *GR-axr2-1* were selected for further analysis.

### Bacterial strains and culture conditions

*Pseudomonas syringae* strain *Pto*DC3000 wild-type was used in this study, and was grown on Nutrient Yeast Glycerol medium [NYG; 57] or in a modified Hrp/Hrc Derepressing Media (HDM) containing 50 mM fructose and 20 μM citrate (30) at 28°C with 100 μg ml^-1^ rifampicin, plus other antibiotics as needed (25 μg ml^-1^ kanamycin, 16 μg ml^-1^ tetracycline). When grown in liquid medium, cultures were shaken at 200 rpm.

### Auxin-responsive gene expression in Arabidopsis plants

To monitor auxin-responsiveness in mature *GR-axr2-1* transgenic and WT Col-0 plants, plants were sprayed with 0.1% ethanol (mock) or 10 mM Dexamethasone (DEX; Sigma-Aldrich, St. Louis, MO, USA) in 0.1% ethanol 24 hours prior to auxin treatment. Fully expanded leaves were infiltrated with a needle-less syringe containing 1 µM of the synthetic auxin Naphthaleneacetic acid (NAA; Sigma-Aldrich, St. Louis, MO, USA) dissolved in 0.01 % Dimethyl sulfoxide (DMSO) or 0.01 % DMSO for mock treatment (0 µM). Plants were allowed to stand at room temperature and the leaves were harvested after 3h, flash-frozen in liquid nitrogen and stored at −80 °C. The frozen samples were ground using a bead beater machine (RETSCH, Newtown, PA, USA) followed by total RNA extraction using the RNeasy Plant Mini Kit (Qiagen, Germantown, MD, USA) according to the manufacturer’s instructions. Residual genomic DNA was digested during RNA purification using an on-column DNAse I treatment (Qiagen, Germantown, MD, USA). The purified RNA was reverse-transcribed to synthesize first strand cDNA using SuperScript™ III Reverse Transcriptase (Thermo Scientific, Waltham, MA, USA) or the Revertaid™ premium first strand cDNA synthesis kit (Thermo Scientific, Waltham, MA, USA). Negative control reactions lacking reverse transcriptase were run in parallel to verify that there was no contamination from genomic DNA. Real time PCR (qPCR) was used to monitor the expression of the auxin-responsive genes *GH3.3* (AT2G23170) and *IAA19* (AT3G15540) (32, 58) using SYBR® Green JumpStart™ Taq ReadyMix™(Sigma-Aldrich, St. Louis, MO, USA) and ig™ SYBR® Green qPCR 2x master mix (Intact Genomic, St Louis, MO, USA) on a CFX Connect real-time PCR detection system (Bio-Rad, Hercules, CA, USA). In each experiment, gene expression analysis was performed on three biological replicates with three technical replicates for each biological replicate.

The cycling conditions were as follows: initial denaturation for 15 min at 95 °C, followed by 40 cycles of 95 °C for 5 sec and 58 °C for 30 sec, with camera capture at the end of each cycle, then 72 °C extension for 30 sec. To confirm the specificity of all amplifications a melt curve was generated after the 40^th^ cycle, using the following parameters: 65 °C for 5 sec, 95 °C for 5 min, then a slow ramp (0.5 °C for 5 s), with camera capture. The relative expression was determined using a relative quantification method of Pfaffl (59, 60). RT-qPCR data were normalized using *Protein Phosphatase 2A subunit A3* [*PP2AA3*, AT1G13320; 61] and *polyubiquitin* 10 [*UBQ10*, AT4G05320; 61] as reference genes. The mock-treated sample was used as a calibrator of relative expression. Auxin-responsive gene expression was carried out in three independent experiments for WT Col-0 (treated with Dex) and in two independent experiments for *GR-axr2-1* plants (treated with Dex or mock-treated) and *GR-axr2-1 sid2* plants treated with Dex. All primers used in this study are described in Supplementary Table 1.

### Pathogen inoculation and *in planta* bacterial growth

*A. thaliana* Col-0 (WT), mutant and transgenic plants were inoculated at approximately 4-5 weeks of age. Bacterial solutions containing ∼10^6^ *Pto*DC3000 cells ml^-1^ in 10 mM MgCl_2_ prepared from freshly-growing bacterial cultures were injected into leaves using a 1 ml needle-less syringe. To quantify bacterial growth in the plant, whole leaves were sampled 2-3 hours after inoculation (day 0) and 3 or 4 days after inoculation, weighed to determine leaf mass, ground in 10 mM MgCl_2_ and then plated in serial dilutions on NYG media with rifampicin. Four to eight leaves were sampled per treatment, depending on the experiment and time point. Following incubation at 28 °C for 48 h, colonies were counted to determine the number of bacteria in the leaves. For experiments involving Dex treatment, plants were sprayed with 0.1 % Ethanol (mock) or 10 µM DEX suspended in 0.1 % ethanol 24 hours prior to inoculation.

### Hormone quantification in plant tissue

To quantify the levels of free IAA and SA, WT Col-0 and *tir1-1 afb1-3 afb4-8 afb5-5* (4x) mutant plants were inoculated with *Pto*DC3000 (10^6^ cfu ml^-1^) or treated with 10 mM MgCl_2_ (mock). 24 hours after inoculation, approximately 100 mg of leaves were sampled for each of three biological replicates per treatment, flash frozen in liquid nitrogen, and stored at −80°C. Subsequently, free IAA and SA were quantified by LC-MS/MS as described in supplemental information.

### Monitoring bacterial gene expression and the effect of IAA in culture

To monitor the effect of IAA on bacterial gene expression in culture, triplicate cultures of 10 mL of NYG broth were inoculated with *Pto*DC3000, and incubated for several hours at 28°C until they reached an OD_600_ of ∼0.07-0.10 as measured by a BioTek PowerWave XS2 96 well plate reader. As a control, a 1 mL sample of bacterial cells was removed from each culture (NYG), treated with 2 mL of RNAprotect bacteria reagent (Qiagen, Germantown, MD, USA) following the manufacturer’s instructions and the samples flash-frozen in liquid nitrogen and stored at −80 °C until further use. The remaining cultures were then transferred to a modified HDM media supplemented with 20 µM citrate (30) to induce expression of T3SS-related genes, containing either IAA or a buffer control. This was accomplished by collecting the cells from the initial NYG cultures by centrifugation at room temperature at 5000 g for 5 min, and resuspending each cell pellet in 10 mL fresh media (HDM), which was then split into two 5 mL aliquots and immediately treated with 100 µM IAA (in 0.1 % DMSO) or 0.1 % DMSO (no IAA control). At 1.5 hours after treatment, 1 mL of each culture was removed and treated with 2 mL of RNAprotect bacteria reagent following, flash-frozen in liquid nitrogen and stored at −80 °C. Bacterial growth was monitored in the cultures prior to and for ∼12 hours after the treatment.

For each *Pto*DC3000 sample, RNA was extracted using the RNeasy RNA Isolation Kit (Qiagen, Germantown, MD, USA). Samples stored at −80 °C were thawed, the cells were lysed enzymatically by treatment with 0.1 mL lysozyme (1 mg/mL in TE buffer), and RNA was extracted following the manufacturer’s instructions using the RNase-free DNase I Set for on-column DNase treatment (Qiagen, Germantown, MD, USA).

For each sample, approximately 1µg of purified RNA was used for cDNA synthesis using SuperScript III (Thermo Scientific, Waltham, MA, USA) and random hexamers as primers (Integrated DNA Technologies, Coralville, IA, USA). Control reactions lacking reverse transcriptase (RT) were included to check the samples for genomic DNA contamination. The products from the cDNA synthesis reactions were diluted into 30 µL of 10 mM Tris buffer (pH 8) and stored at −4 °C. In order to verify the quality of the cDNA and to make sure that there was no genomic DNA contamination, PCR reactions were performed on all samples using primers for 16S rRNA (Supplemental Table 1), using the following cycling conditions: 5 min at 95 °C, followed by 25 cycles of 95 °C for 30 sec, 58 °C for 30 sec, and 68 °C for 3 min. The amplification product of this reaction was visualized as a ∼1.5 kb band on an agarose gel. Only cDNA samples that exhibited no DNA contamination in the control reactions lacking RT were used for RT-qPCR. Expression of the following virulence-related genes was monitored: *avrPto* (*PSPTO_4001*), *hrpL* (*PSPTO_1404*), *cmaA* (*PSPTO_4709*), *tvrR* (*PSPTO_3576*), *hcp1* (*PSPTO_2539*) and *PSPTO_5415,* which is predicted to encode an Rhs element Vgr protein (www.pseudomonas.com). To monitor bacterial gene expression, qPCR was performed on cDNA samples using PowerUp SYBR Green Master Mix (Thermo Scientific, Waltham, MA, USA). Reactions were set up in a 20µL final volume, and qPCR was performed on a CFX Connect Real-Time PCR Detection System (Bio-Rad, Hercules, CA, USA). The cycling conditions were as follows: 2 min at 94 °C, followed by 49 cycles of 95 °C for 15 sec and 58 °C for 30 sec, with camera capture at the end of each cycle. The specificity of all amplifications was confirmed by generating a melt curve after the 40^th^ cycle, using these parameters: 65 °C for 5 sec, 95 °C for 5 min, then a slow ramp (0.5 °C for 5 sec), with camera capture. Bacterial gene expression was calculated using the comparative Ct method for relative quantification (62). Each biological replicate was tested in technical triplicates, and the Cq for each biological replicate (average of three technical replicates) was normalized to the geometric mean of two internal reference genes, *gyrB* (*PSPTO_0004*) and *rpoD* (*PSPTO_0537*). Bacterial gene expression from the NYG samples collected prior to transfer of the culture to HDM was used as a calibrator of relative expression. All primers used in this study are described in Supplementary Table 1.

### Monitoring bacterial gene expression *in planta*

*Arabidopsis* WT Col-0 and *tir1-1 afb1-3 afb4-8 afb5-5* mutant plants were inoculated at approximately 4-5 weeks of age. Whole leaves were syringe-infiltrated with *Pto*DC3000 (10^6^ cfu ml^-1^) in 10 mM MgCl_2_ prepared from freshly-growing bacterial cultures or treated with 10 mM MgCl_2_ (mock treatment for *in planta* gene expression). Approximately 100 mg of leaves were collected for RNA isolation at 24 and 48 h after inoculation, frozen immediately in liquid nitrogen and stored at −80 °C. A combination of protocols from the RNAprotect bacteria reagent kit (Qiagen, Germantown, MD, USA) and RNeasy Plant Mini kit (Qiagen, Germantown, MD, USA) was used to isolate and enrich for bacterial RNAs from the samples. The frozen leaves were ground into a fine powder using a bead beater machine (RETSCH, Newtown, PA, USA), followed by addition of 1.0 mL of RNAprotect bacteria reagent to each sample. The standard protocol for enzymatic lysis of bacteria from the RNAprotect bacteria reagent handbook (Protocol 1 up to step 9, https://www.qiagen.com) was performed. Following the addition of the RLT lysis buffer, the protocol from the RNeasy Plant Mini kit was followed. The total RNA obtained was a mix of bacterial and plant RNAs. We used the concentration of *Pto*DC3000–specific 16S rRNA (PSPTO_r01) as an in-sample proxy for bacterial concentration within the mixed RNA samples as described recently by Smith et al (36). In this approach, the concentrations of bacterial RNA in the total RNA isolated from infected plant tissue were standardized using *Pto*DC3000–specific 16S ribosomal RNA primers. For comparison, 1.0 ml of the initial inoculum was collected by centrifugation and total RNA extracted following the protocol described above for bacteria grown in culture. The extracted RNA was used for cDNA synthesis using the Revertaid™ premium first strand cDNA synthesis kit (Thermo Scientific, Waltham, MA, USA) and random hexamers with the subsequent steps exactly as mentioned above for the bacterial cultures. qPCR was then used to monitor the expression of *avrPto* (*PSPTO_4001*), *hrpL* (*PSPTO_1404*), *cmaA* (PSPTO_4709) and *tvrR* (PSPTO_3576) using ig™ SYBR® Green qPCR 2x master mix (Intact Genomic, St Louis, MO, USA) on a CFX Connect real-time PCR detection system (Bio-Rad, Hercules, CA, USA). The cycling conditions were: 15 min at 95 °C, followed by 40 cycles of 95 °C for 5 sec and 58 °C for 30 sec, with camera capture at the end of each cycle. A melt curve was generated after the 40^th^ cycle, using these parameters: 65 °C for 5 sec, 95 °C for 5 min, then a slow ramp (0.5 °C for 5 sec), with camera capture. In each experiment, gene expression analysis was performed on three biological replicates with three technical replicates for each. The relative expression was determined using a relative quantitation method of Pfaffl as described previously (59, 60). RT-qPCR data were normalized to *recA* (*PSPTO_4033*) and *rpoD* (*PSPTO_0537*) used as reference genes. The stability of *recA* and *rpoD* expression in planta at 24 and 48 h after inoculation was confirmed as described by Smith et al. (36), using a synthetic double-stranded DNA gBlock (Integrated DNA Technologies, Coralville, IA, USA) and *Pto*DC3000–specific *16S rRNA* primers. The bacterial gene expression in the inoculum sample was used as a calibrator of relative expression. We also monitored the expression of *PATHOGENESIS RELATED PROTEIN 1 (PR1)* and *IAA-19* in these samples, in leaves harvested 24 h after inoculation, following the protocol mentioned above for auxin responsive genes.

### Statistical analysis

Data sets were statistically compared with the statistical analysis software GraphPad Prism 8.0 (GraphPad software, San Diego, CA, USA) using one-way analysis of variation (ANOVA), followed by the Tukey’s post-test. The confidence level of all analyses was set at 95 %, and values with p < 0.05 were considered significant.

## ACKNOWLEDGMENTS

We are grateful to Zipeng (Alex) Li, Cynthia Holland, Anne Zimmerman, Sarah DeCou and Saryu Sanghani for technical help with experiments. We thank Catherine Perrot-Rechenmann and Lucia Strader for helpful discussion and Rebecca Bart for comments on the manuscript. We also thank Brad Evans, Jonathan Mattingly, Jia Li and Sophie Alvarez at the Proteomics and Mass Spectrometry Facility of the Donald Danforth Plant Science Center (St. Louis, MO) for hormone analysis. Their analytical methods are based upon work supported by the National Science Foundation under Grant No. DBI-1427621 for acquisition of the QTRAP LC-MS/MS. This work was supported by grants from the National Science Foundation (IOS-1645908) awarded to BNK and National Institutes of Health (GM43644) awarded to M.E.

