## Supplemental Information & Figures for "Dual role of auxin in regulating plant defense and bacterial virulence gene expression during *Pseudomonas syringae Pto*DC3000 pathogenesis"

**Plant Hormone Quantification: IAA and SA Extraction and Quantification**

Materials

Indole-3-acetic acid (IAA) and salicylic acid (SA) were purchased from Sigma-Aldrich (St. Louis, MO) and Acros Organics (Geel, Belgium), respectively. Isotope-labeled internal standards $d_4$-salicylic acid ($d_4$-SA) and indole-2,4,5,6,7-$d_5$-3-acetic acid ($d_5$-IAA) were sourced from CDN Isotopes (Pointe-Claire, Quebec). Methanol and acetonitrile (HPLC-grade) were sourced from J.T. Baker (Avantor Performance Materials) and LC-MS grade water was purchased from Honeywell Research Chemicals. Individual stock solutions of unlabeled and labeled compounds were prepared in 50% methanol and stored at -80 °C. Standard solutions were prepared fresh in 30% methanol in the linear concentration ranges of 32 pM to 10 µM. An internal standard solution was prepared in 30% methanol containing 2.5 µM $d_5$-IAA and $d_4$-SA.

Plant Hormone Extraction

Frozen plant material was extracted with 900 µL of extraction solvent (ice-cold acetonitrile/methanol; 1:1 v:v) while samples were kept cold on ice. Ten (10) microliters of the internal standard mixture and two stainless steel 5 mm beads were added to each sample tube followed by brief mixing by vortexing. Samples were placed in pre-cooled (-80 °C) Tissue Lyser II racks and homogenized for 2 min at 15 Hz. Samples were centrifuged at full speed for 5 min at 4 °C, then the supernatant was transferred to a new 2 mL tube. Samples were re-extracted with another 900 µL of extraction solvent, and then homogenized again for 2 min at 15 Hz. The samples were then centrifuged and supernatant transferred as previously described. The extraction solvent was removed under reduced pressure with a speed-vac until completely dry. Samples were reconstituted in 30 % MeOH (200 µL) and mixed thoroughly for 30 min at 4 °C. Finally, samples were filtered through 0.8 µm PES spin-filters and 40 µL of clarified supernatant was transferred to a 96-well microplate. Two (2) µL of sample was injected onto the column.

LC-MS/MS Instrumentation

Clarified samples were analyzed on an Eksigent ekspert™ microLC 200 coupled to a Sciex 6500 QTrap® (Framingham, MA) operated with polarity-switching electrospray ionization. The LC separation was achieved using a Waters (Milford, MA) Acquity UPLC® BEH C18 1.0 × 100 mm, 1.7 µm column kept at 50 °C with a flow rate of 15 µL/min while the autosampler was set at 8 °C. The mobile phases were 0.1 % acetic acid and 3:1 acetonitrile:methanol containing 0.1 % acetic acid running a gradient of 20 % B for 4 minutes ramping to 70 % B at 7 minutes, increasing to 95 % B at 7.5 minutes, holding for 5.5 minutes, then re-equilibrate at initial conditions at 13.5 minutes for 10 minutes (total runtime is 23.5 minutes). Data analysis was completed using MultiQuant 3.0.2 (AB Sciex) by normalizing the peak areas of the unlabeled analytes relative to the peak areas of the labeled internal standards. Calibration curves were linear (*r* values = > 0.99) within the ranges provided above applying a 1/x weighting scheme.

Compound-dependent parameters

| Compound | MRM Transition | Retention time (min) | DP (V) | EP (V) | CE (V) | CXP (V) |
|---|---|---|---|---|---|---|
| IAA | 176.0 → 130.0 | 10.24 | 23 | 7 | 40 | 19 |
| $d_5$-IAA | 181.0 → 134.0 | 10.22 | 23 | 7 | 25 | 19 |
| SA | 137.0 → 93.0 | 10.20 | -22 | -13 | -20 | -10 |
| $d_4$-SA | 141.0 → 97.0 | 10.18 | -41 | -9 | -23 | -11 |

Source settings

Ionspray voltage: polarity switching between -4500 V in –ve to 4500 V in +ve

Curtain gas: 15

Gas 1: 35

Gas 2: 35

### Supplemental Table 1 Primers used in this study

| Gene | Forward primer (5'-3') | Reverse primer (5'-3') | Reference for primers |
|---|---|---|---|
| | | **RT-qPCR** | |
| *GH3.3* *AT2G23170* | TGTGTGAGTTTCACGCCAAT | CAAAGGAGGGACAGAGTGGA | (32) |
| *IAA19* *AT3G15540* | GAGATGTGGCAGAGAAGATG | TTCCTCAAATAAGGCACACC | (32) |
| *PR1* *AT2G14610* | GGAGCTACGCAGAACAACTAAGA | CCCACGAGGATCATAGTTGCAACTGA | (32) |
| *PP2AA3* *AT1G13320* | AACGTGGCCAAAATGATG | AACCGCTTGGTCGACTATCG | (61) |
| *UBQ10* At4g05320 | CGTTAAGACGTTGACTGGGAAAACT | GCTTTCACGTTATCAATGGTGTCA | (61) |
| *avrPto* *PSPTO_4001* | ATGACGGGAGCGTCAGGAATCAAT | ATCCGTTCGGGTTCATAGTCGCAA | (30) |
| *HrpL* *PSPTO_1404* | TCAGGAAAGCTGGGAAGAC-GAAGT | ATGTTCGACGGCAGGCAATCAATG | (30) |
| *mqo* *PSPTO_1136* | GCGGCTGATGGCTCCATCGAC | CGGGACCGGATTGATGAACGAC | This work |
| *cmaA* *PSPTO_4709* | CCGTGATGTTTACCTCTGGCAC | GGACGAGTGATGTACGTAGCTGC | This work |
| *hcp1* *PSPTO_2539* | GGTCGACGCAGGCATAACGC | CTCCTTGCCGTCGTTAGTGCG | This work |
| *tvrR* *PSPTO_3576* | GGCTCGCAACGGCCCATCTG | CATGCGGTAGACGGCCAGCG | This work |
| *PSPTO_5415* | GCCAGGAAGGGCATGTGCTG | AATCCCTTGATGACCGGCACG | This work |
| *16S rRNA* *PSPTO_r01* | TAATGGCTCACCAAGGCGACG | TGGCTGGATCAGGCTTTCGC | This work |
| *rpoD* *PSPTO_0537* | GAAGTTGACGAAAGCTGGACCG | CGACGGTTGATGTCCTTGATCTC | This work |
| *gyrB* *PSPTO_0004* | CTTCAGCTGGGACATTCTGGC | AACCGCCTTCGTACTTGAACAG | This work |
| *recA* *PSPTO_4033* | TAGAACTTCAGCGCGTTACC | GCCAACTGCCTGGTTATCT | (36) |
| | **Genotyping (PCR) and Cloning** | | |
| *sid2-2 sm108F* | TTCTTCATGCAGGGGAGGAG | AAGCAAAATGTTTGAGTCAGCA | (32) |
| *WT- Sm30F* | CAACCACCTGGTGCACCAGC | | |
| *GR-axr2-1* | GR: GCCATCGTCAAAAGGGAAGG | | This work |
| *axr2* | | TGACTCTAACTCGGTAAGGTTCAT | This work |
| *TopoGW_GR__F1* | CACCATGATTCAGCAAGCCACTGC | | This work |
| *GR_AXR2_R1* | | TGAGGTTCATAAGTTGGCCGAT CATTTTTTGATGAAACAGAAGCT | This work |
| *GR_AXR2_F1* | AGCTTCTGTTTCATCAAAAAATG ATCGGCCAACTTATGAACCTCA | | This work |
| *AXR2_V1_R1* | | TCAAGATCTGTTCTTGCAGTACT | This work |

### Supplemental Figures

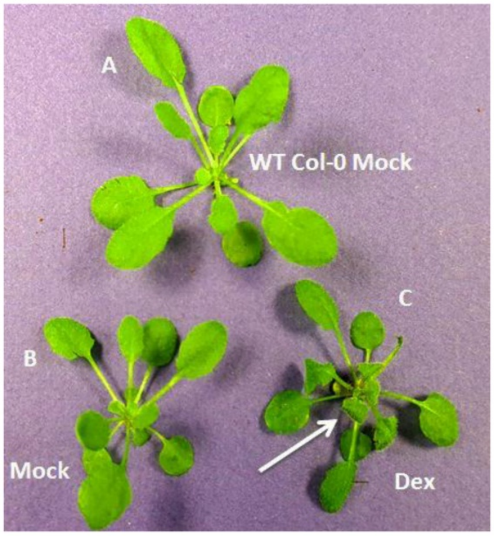


**Supplemental Fig. 1. Dex treatment of *GR-axr2*-1 transgenic plants results in abnormal leaf morphology. (A)** WT Col-0; **(B)** and **(C)** *GR-axr2*-1 transgenic plants, 1 day after spraying with 0.1 % Ethanol (Mock) or 10 mM Dexamethasone in 0.1 % Ethanol (Dex). Arrow indicates curled leaves that did not expand normally.

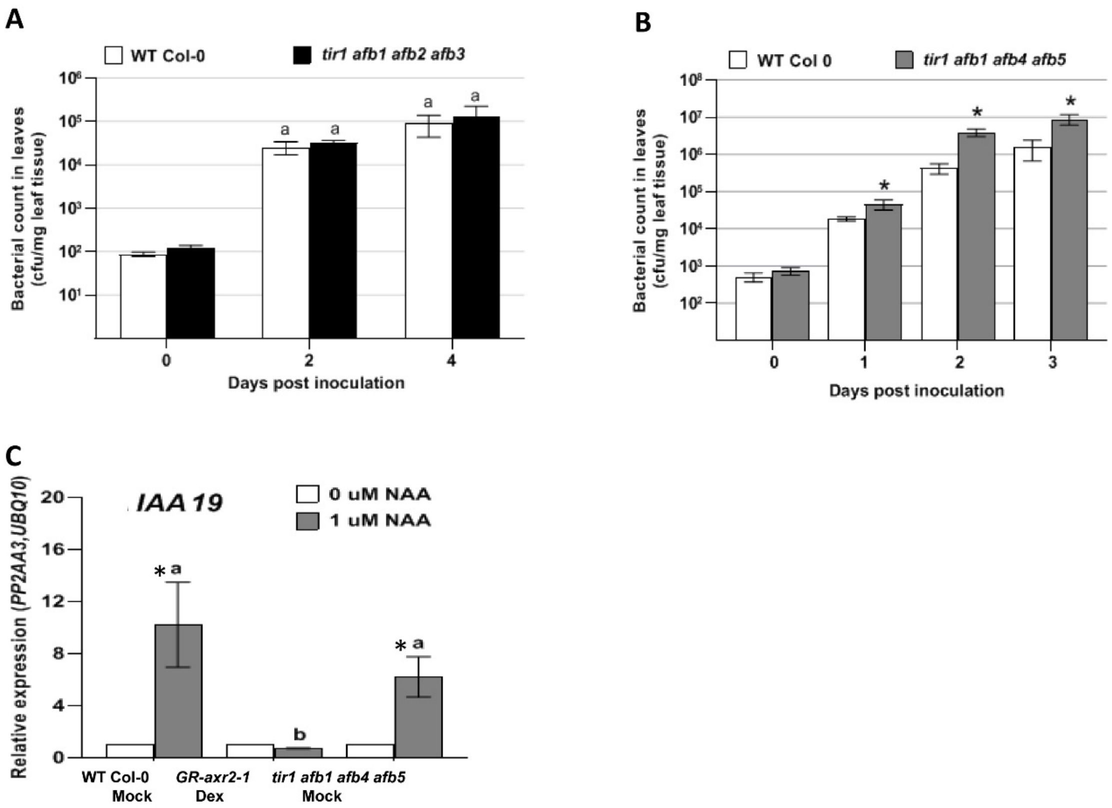


**Supplemental Fig. 2. Growth of *Pto*DC3000 and expression of auxin-responsive genes *IAA19* in auxin receptor mutants.**

**(A)** Growth of *Pto*DC3000 in the *tir1 afb1 afb2 afb3* mutant. Similar results were observed in 3 additional independent experiments. **(B)** Growth of *Pto*DC3000 in WT Col-0 and *tir1 afb1 afb4 afb5* plants. Values are an average ± SEM for data from 4 biological replicates (leaves) for day 0 and 8 replicates for days 1, 2, and 3. Results were analyzed using ANOVA, followed by a Tukey's post test, and asterisks (*) indicates significant differences between WT Col-0 and *tir1 afb1 afb4 afb5* (4X) mutant plants with P < 0.05 at each time point. Plants were inoculated with a bacterial suspension of $10^6$ cfu $ml^{-1}$. Samples from this experiment were used for gene expression experiments shown in Fig. 5. **(C)** Expression of the auxin-responsive gene *IAA19* after NAA treatment. Values are an average ± SEM for 3 biological replicates for Col-0 WT Mock, *GR-axr2-1* and *tir1 afb1 afb4 afb5*. Samples indicated by different lower-case letters are significantly different (p<0.05). Similar results were obtained in a second independent experiment. * indicates significant difference between treatment (0 and 1 µM NAA) with p<0.05. Results were analyzed using ANOVA, followed by a Tukey's post test. Samples indicated by different letters are significantly different at p<0.05.

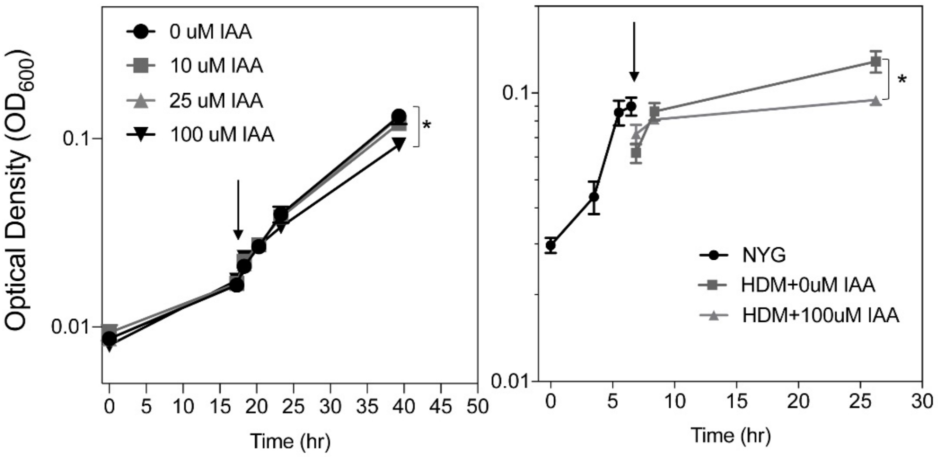


**Supplemental Figure 3. Impact of IAA on growth of *Pto*DC3000 in culture**
**(A)** Growth of *Pto*DC3000 in Hrp De-repressing Medium (HDM) containing the indicated concentrations of IAA. IAA was added to the cultures at mid-log (arrow), and growth was monitored by quantifying cell density ($OD_{600}$) over ~ 24 hours. **(B**) *Pto*DC3000 was grown in NYG to an OD of ~0.1, cells were collected via centrifugation, resuspended in NYG or HDM (arrow) containing the indicated concentrations of IAA and grown for ~24 hours. Growth was monitored by quantifying cell density ($OD_{600}$). Cells were collected at 1.5 hours after transfer to HDM for RNA isolation. Cell cultures were grown in triplicate for both experiments. Data points are the average of 3 biological replicates per treatment and error bars represent S.E.M. A student's t-test was performed to compare the effect of the IAA treatments on bacterial growth to Mock treatment with DMSO. * $p<0.05$ . The data shown in panel B are from cultures used to monitor the effect of IAA on PtoDC3000 gene expression shown in figure 4.
